# Introns structure patterns of variation in nucleotide composition in *Arabidopsis thaliana* and rice protein-coding genes

**DOI:** 10.1101/010819

**Authors:** Adrienne Ressayre, Sylvain Glémin, Pierre Montalent, Laurana Serre-Giardi, Christine Dillmann, Johann Joets

## Abstract

Plant genomes are large, intron-rich and present a wide range of variation in coding region *G* + *C* content. Concerning coding regions, a sort of syndrome can be described in plants: the increase in *G* + *C* content is associated with both the increase in heterogeneity among genes within a genome and the increase in variation across genes. Taking advantage of the large number of genes composing plant genomes and the wide range of variation in gene intron number, we performed a comprehensive survey of the patterns of variation in *G* + *C* content at different scales from the nucleotide level to the genome scale in two species *Arabidopsis thaliana* and *Oryza sativa*, comparing the patterns in genes with different intron numbers. In both species, we observed a pervasive effect of gene intron number and location along genes on *G* + *C* content, codon and amino acid frequencies suggesting that in both species, introns have a barrier effect structuring *G* + *C* content along genes. In external gene regions (located upstream first or downstream last intron), species-specific factors are shaping *G* + *C* content while in internal gene regions (surrounded by introns), *G* + *C* content is constrained to remain within a range common to both species. In rice, introns appear as a major determinant of gene *G* + *C* content while in *A. thaliana* introns have a weaker but significant effect. The structuring effect of introns in both species is susceptible to explain the *G* + *C* content syndrome observed in plants.

## INTRODUCTION

In Eukaryotes, protein-coding genes are formed of an alternation of coding and non-coding regions (Fig. 1). Non-coding regions are the 5’ and 3’ Untranslated Terminal Regions (UTRs) and the spliceosomal introns. During transcription and messenger RNA (mRNA) maturation, introns are excised by a large cellular machinery, the spliceosome (Reddy, 2007). Coding sequences and UTRs are then spliced to form the mature mRNA composed of the concatenation of the coding sequences surrounded at each extremity by the 5’ and 3’ UTR. Although introns bear no information regarding mRNA or protein sequences, introns or splicing processes are implicated in a wide range of critical processes regarding gene expression (Lynch, 2002; Maniatis and Reed, 2002; Moore and Proudfoot, 2009; Carmel and Chorev, 2012). Implication of the splicing processes are observed throughout gene expression from transcription initiation and 5’-capping to poly-adenylation, export from the nucleus and even the first round of translation. Alternative splicing not only contributes to the expansion of the proteome but also affects gene expression (Kelemen et al., 2012; Syed et al., 2012). In both plants and animals, intron presence is generally associated with an increase in protein production mediated through many different mechanisms ranging from the increase of transcription and translation rates to the improvement of mRNA stability and folding (Lorkovic et al., 2000; Le Hir et al., 2003; Ren et al., 2006; Karve et al., 2011; Skoko et al., 2011; Carmel and Chorev, 2012; Moabbi et al., 2012). Moreover, an unknown proportion of introns contains non-coding RNAs, ORFs or signaling sequences required for gene expression (Lynch, 2002; Karve et al., 2011; Skoko et al., 2011; Carmel and Chorev, 2012). All these functional aspects are associated with a range of selective pressures unrelated to protein sequence (Warnecke et al., 2009; Weatheritt and Babu, 2013) but that indirectly affects molecular rates of protein evolution. Correct splicing or alternative splicing is required for the production of functional transcript and the canonical splicing motifs (5’ and 3’ splice sites) or the enhancer motifs located near splice junctions are under selection and affect codon usage in neighboring coding sequences in a variety of eukaryotes (Comeron and Guthrie, 2005; Larracuente et al., 2008; Ke et al., 2008; Warnecke and Hurst, 2007; Parmley et al., 2007; Denisov et al., 2014; Falanga et al., 2014).

**Figure 1:**
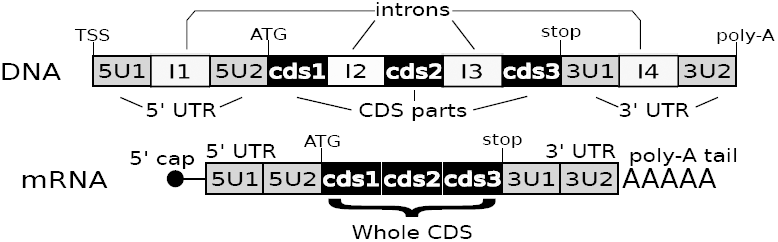
Protein-coding gene structure. A gene is composed of a variable number of regions that can be of 4 types: 5’ UnTranslated Regions (UTR), coding regions (CDS), intron and 3’ UTR. Introns are non-coding regions interrupting any of the other types of regions. They are excised and the remaining regions are spliced to produce the mature mRNA. 5’ and 3’ UTRs occupy 5’ and 3’ gene and mRNA extremities. Coding sequences encode for the amino acid sequence of the protein. Along a gene, each region was denoted according to its type and its rank position from the 5’ end towards the 3’ end. In the example shown in the figure, the gene has four introns, one intron in each UTRs and two introns in the coding region. Its decomposition in elements is provided by the list (5U1, I1, 5U2, cds1, I2, cds2, I3, cds3, 3U1, I4, 3U2), the whole CDS consisting in the three concatenated CDS parts (cds1cds2cds3). CDS will be used to refer to the complete coding sequence of a gene regardless of whether it is interrupted or not by introns. TSS is for transcription start site.

Beyond selection on canonical splice sites or enhancer motifs, a range of observations suggests that introns might have an influence on coding region nucleotide (nt) composition (*G*+*C* content). In both plants and animals, the differences between *G*+*C*-rich coding regions and *G*+*C*-poor introns are linked with splicing efficiency (Goodall and Filipowicz, 1989, 1991; Carle-Urioste et al., 1997; Amit et al., 2012). The intron-exon architecture of the genes also overlaps with chromatin organization, nucleosomes occupying preferentially exons while linkers are mainly formed by introns (Andersson et al., 2009; Chodavarapu et al., 2010; Amit et al., 2012), and nucleosome occupancy being mainly determined by sequence *G* + *C* content (Tillo and Hughes, 2009). Furthermore intron number, intron location and/or intron length are recurrently reported to affect *G* + *C* content of both introns and exons (Carels and Bernardi, 2000; Wang et al., 2004; Guo et al., 2007b; Zhu et al., 2009; Amit et al., 2012). Recently, two studies in plants revealed the existence of a close association between patterns of variation in *G* + *C* content in coding regions, introns and recombination rates (Choi et al., 2013; Hellsten et al., 2013), providing a potential mechanism to explain how the intron-exon architecture of genes can influence nt composition through GC-biased gene conversion, a process associated with recombination in several eukaryotes that favors the transmission of G and C alleles at meiosis (Duret and Galtier, 2009; Glémin et al., 2014). All these reports collectively suggest that at least in plants, introns could influence patterns of variation in *G* + *C* content of genes and we decided to address this question in two plant genomes with contrasted *G* + *C* content, *A. thaliana* and *Oryza sativa* (rice).

In plants, coding sequence *G* + *C* content vary continuously from 40 to 60% depending on the species, with *A. thaliana* and rice being representative of *G* + *C*-poor and *G* + *C*-rich species respectively (Serres-Giardi et al., 2012). Like *A. thaliana*, *G* + *C*-poor plant genomes display low level of variations among genes (fig. 2A). Throughout plants, the increase in genome-wide *G* + *C* content is associated with an increase in *G* + *C* content variability among genes leading in *G* + *C*-rich genomes to a bimodal distribution of coding sequence *G* + *C* content as observed in rice (fig. 2B). In both species, the association between *G* + *C* content and gene intron number is obvious, *G* + *C*-rich genes displaying low intron number while *G* + *C*-poor genes tend to be intron-rich. Within plant genomes (including *A. thaliana* and rice), coding region *G* +*C* content also varies along genes, following a 5’ to 3’ decreasing *G* + *C* content gradients scaled with genome-wide *G* + *C* content being described (Wong et al., 2002; Serres-Giardi et al., 2012). Gradients along genes of varying amplitudes could explain why *G* + *C*-poor plant genomes display little *G* + *C* content variation among genes while *G* + *C*-rich genome are highly heterogeneous regarding *G* + *C* content (Wong et al., 2002; Glémin et al., 2014). If gradients along genes are steep, simple variations in coding region length can cause large variations in *G* + *C* content. If in addition, introns contribute to *G* + *C* content gradients for example because they interfere with GC-biased gene conversion, intron number could explain at least a part of the variation existing among genes within plant genomes. To test for this hypothesis, we re-analyzed the genomes of *A. thaliana* and rice focusing our analysis on the consequences of intron presence on *G* + *C* content at different scales and investigating the effect of changes in intron structure on *G* + *C* content by studying the impact of intron insertion outside coding regions within the 5’ or the 3’ UTRs. Our results converge to show that in both species, introns are intricately associated with *G* + *C* content patterns of variation at any of the studied scales and appear as a major determinant of nt composition.

**Figure 2:**
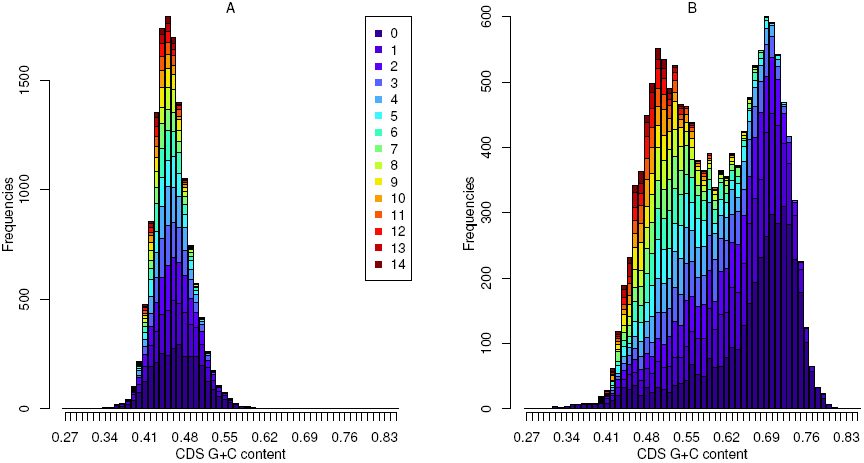
Coding region *G*+*C* content distribution and intron number in *A. thaliana* (A) and *O. sativa* (B) for genes with less than 15 introns inserted in the coding sequences and no intron inserted in the UTRs. The contribution of the intron number class to each of the bars is indicated by the proportion of the bar of the relevant color (legend in panel A). A. *A. thaliana* distribution is unimodal. B. *O. sativa* distribution is bimodal and almost symmetrical. In both species, *G* + *C*-rich bars are mainly composed of genes with a low intron number, while genes with a high intron number are mainly concentrated in the *G* + *C*-poor classes.

## RESULTS

In each species, we filtered genomic data removing genes with missing 5’ or 3’ UTRs, not supported by full-length cDNA or having more than a single intron present either in their 5’ or 3’ UTR. The remaining set of genes were further divided into three different subsets of genes. The first subset comprised genes with no intron located either in the 5’ nor the 3’ UTR, the second subset genes with a single intron located within the 5’ UTR and no intron located in the 3’ UTR and the third subset genes with an intron located within the 3’ UTR and no intron located in the 5’ UTR. The first subset representing more than 75% of the genes was used to describe patterns of variation linked with intron presence and the two other gene subsets were used to investigate the consequences of intron insertion on *G* + *C* content of both introns and coding regions. Coding part of exons and UTR were analyzed separately because they exhibit different *G* + *C* patterns (see Materials and Methods). To avoid terminology confusion we discarded the term “exons” and used the term “CDS parts” for coding parts of mature mRNA. In both species, coding region *G* + *C* content varying with intron number (fig. 2), all gene subsets were further sorted according to intron number. In the intron-free UTR subsets, gene intron number distribution is fairly similar between the two species, except for the genes with low intron number that are in excess in rice when compared to *A. thaliana* (for additional information see Supplementary Table S1).

### Introns organize patterns of *G* + *C* content variation along genes into gradients

Patterns of variation in *G* + *C* content along genes are fairly complex, three intermingled levels of structuring being described, (i) systematic differences larger than 10% between introns and coding regions, transitions between regions being abrupt (Goodall and Filipowicz, 1989, 1991; Carle-Urioste et al., 1997), (ii) systematic differences between codon position within coding regions (Wong et al., 2002) and (iii) decreasing 5’ to 3’ *G* + *C* gradients of varying amplitudes in both species (Wong et al., 2002). Gradients have been also observed when the units of observation were the CDS parts or the introns (Guo et al., 2007a; Zhu et al., 2009). All these patterns were usually described without taking into account either gene intron number or gene precise architecture.

To investigate these two aspects, we sorted genes according to intron number and first investigated within each class of genes, patterns of variation at junctions between coding regions and introns as described in Goodall and Filipowicz (1989). We indeed observed that in both species for any intron number and any junction between coding regions and introns, abrupt changes in *G* + *C* content, larger than 10%, take place (examples in both species in supplementary figures S1 and S2). To further characterize patterns of variation associated with introns in coding regions, we then removed intron sequences and aligned concatenated coding regions on their starting methionine only keeping, in addition to nt position, information concerning the rank of the CDS part of the nt.

In both species at nt level and within all intron number classes, large oscillations in *G* + *C* content are observed between neighboring nts (shown for intronless genes, fig. 3AC, and genes with six introns in supplementary figures S3-S4). These oscillations decrease sharply when each codon position is treated separately revealing the existence of three distinct patterns of variation, one per codon position along the coding regions. The codon *G* + *C* content which results from the average of the three codon positions (blue lines in fig. 3AC) occupies an intermediate position and displays on the first hand a smaller range of variation along intronless genes or among rank in intronic genes and on the other hand smaller oscillations between neighboring codons. In addition, we observed that intronless genes display codon *G* + *C* gradients that differ from gradients of all intronic genes and that intron presence is associated with discrete changes in *G* + *C* content between contiguous CDS parts (fig. 3 for codon *G* + *C* content and supplementary figures S3 and S4 for *GC*1, *GC*2 and *GC*3 in genes with 6 introns). As shown in fig. 3D for rice genes with six introns, consecutive CDS parts are overlapping over a range of nt positions because of the wide variation in CDS part lengths among genes. This enables to gauge for differences between different CDS parts at same nt position with respect to the starting methionine. In rice, discrete changes in codon *G* + *C* content are observed between overlapping CDS parts, the continuous gradient observed when neglecting intron cutting up being an artifact caused by the transition of an increasing number of sequences from a given CDS part to the next and not by a progressive change in *G* + *C* content according to codon position. As a result in rice, intron presence is associated with the formation of step-gradient, arrows located in the right side of the panel in fig. 3D indicating that average CDS part *G* + *C* content provides a reasonable approximation for the within CDS part *G* + *C* content variation. Precise information on intron location is required to correctly capture patterns of variation along the genes, an equal-sized bin cutting up consistently underestimating gradient depths for most intron number classes at least in the 5’ regions of the genes (for additional information see supplementary figure S6). In *A. thaliana*, although variation among CDS parts are hidden by nt oscillations within CDS parts (fig.3 B), bin cutting up also underestimates CDS part gradient (see supplementary figure S5 for all intron numbers) at least in the 5’ regions of genes for most intron number classes, indicating that in this species as well, significant changes in *G* + *C* content occur between contiguous CDS parts.

**Figure 3:**
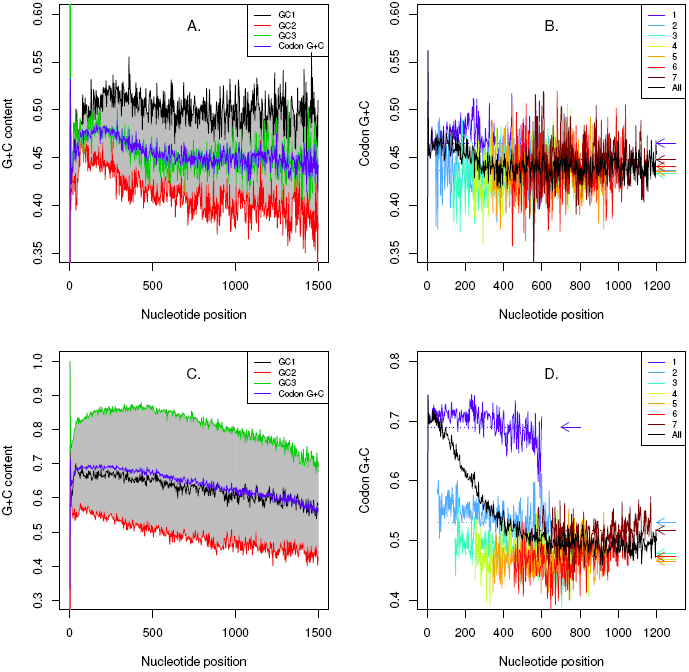
Patterns of variation in coding regions at nucleotide scales. AC. Nt *G* + *C* content according to distance from the start codon at first (black), second (red) and third (green) codon positions within intronless genes (A: *A. thaliana*, C. *O. sativa*). The codon *G* + *C* content (average over the three codon positions, blue) is displayed at the second codon position. Each codon position exhibits its own pattern of variation producing large oscillations when all nts are linked (grey). BD. Codon *G* + *C* content according to distance from the start codon for genes with six introns inserted in the coding regions in *A. thaliana* (B) and *O. sativa* (D). The codon *G* + *C* content (displayed at the second codon position) were computed either without taking into account intron cutting up (black line) or by sub-setting over the genes according to each CDS part rank (blue to brown lines, color legend in the panels). Each CDS part codon *G* + *C* content was plotted for codon positions represented by at least 50 different sequences. In rice (D), a 5’ to 3’ monotonous/continuous/decreasing gradient is observed when intron cutting up is neglected (black line). When intron cutting up is taken into account, the gradient is brocken into a discontinuous step gradient conveniently captured by the average CDS part *G* + *C* content (dotted lines indicated by arrows). In *A. thaliana*, a step gradient is also present (arrows) but hidden by the within CDS part oscillations.

Hence in both species, intron presence is associated with the formation of step gradients in coding regions. In figure 4, we summarized the patterns of variation in *G* + *C* content in both introns and CDS parts by averaging over ranks along genes within intron number classes. Patterns of variation of both CDS parts and introns, and within CDS parts for each codon position, are organized in structured gradients modulated by intron number. These gradients vary in shape, amplitude and regularity according to the types of element (intron or coding region) or the codon position in a species-specific way. However, within a species and for a given element or codon position, gradient shapes tend to be conserved among intron number classes. In both species in the subsample of genes with no intron inserted within UTRs, asymmetric U-shaped gradients in average CDS part *G* + *C* content arise progressively as intron number increases, except for genes with less than two or three introns for which they are truncated (fig. 4A1B1). Within species among intron number classes, the gradients are fairly regular and consistent. The highest *G* + *C* content levels, up to 48% and almost 70% in *G* + *C* content in *Arabidopsis* and rice respectively, are observed in first 5’ CDS part of the genes. Within a few steps the gradients decrease down to 42 − 43% in both species (the third or the fifth intron in *Arabidopsis* and rice respectively for genes with high intron numbers). Then the gradient increases again in both species towards the 3’ end of the CDS and stabilize at intermediate *G* + *C* levels around 44 − 45% in *A. thaliana* and 50% in rice. In both species, the gradients are modulated by intron number, firstly decreasing more as intron number increases until reaching the common lower limit described above nine introns (gradient amplitudes are listed in the supplementary tables S3 for *A. thaliana* and S4 for *O. sativa*), and then enlarging only in width as intron number further increases. As a result in the two species, both CDS part rank and gene intron number affect average CDS part *G* + *C* content, differences among genes within genomes and between the two genomes tending to decrease with the increase in intron number and the rank of the CDS part along the genes. In other words, internal CDS parts of genes with high intron number tend to be similar.

**Figure 4:**
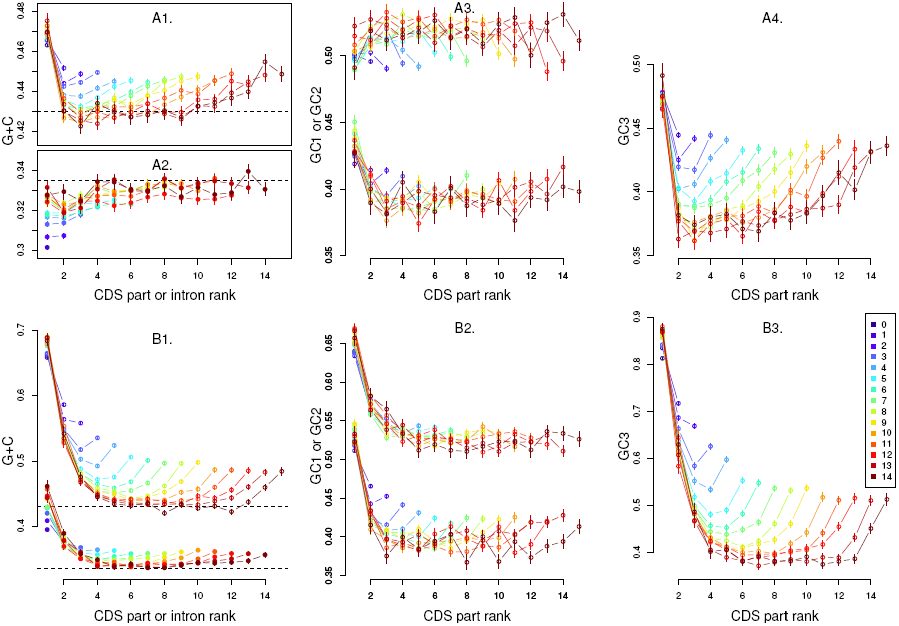
Patterns of variation in CDS part and intron average *G* + *C* content according to rank along genes within intron number classes for genes with less than 15 introns inserted in the coding sequences and no intron inserted in the UTRs. *A. thaliana*: A1. Average CDS part *G* + *C* content. A2. Average Intron *G* + *C* content. A3. Average CDS part *GC*1 (upper groups of lines) and *GC*2 (lower group of lines). A4. Average CDS part *GC*3. *O. sativa*: B1. Average CDS part (upper group of lines) and intron (lower group of lines) *G* + *C* content. B2. Average CDS part *GC*1 (upper groups of lines) and *GC*2 (lower group of lines). B3. Average CDS part *GC*3. In all panels, gene intron number is indicated by the colors (legend shown in panel B3). The upper dashed lines in the B1 panel is placed at the same level as the dashed line in panel A1. Likewise the lower dashed line in the panel B1 is placed at the same level as the one in the panel A2 indicating that both species internal introns tend to reach similar *G* + *C* as intron number increases. Bars on dots represent standard error of means.

In addition, three different CDS part gradients, one for each codon position, are also observed at CDS part level. Except for first codon position in *A. thaliana*, all gradients tend to decrease or to be U-shaped in both species. Compared to the corresponding average CDS part *G* + *C* gradients in each species (fig. 4A1-B1), average CDS part *GC*1 and *GC*2 gradients of both species are shrunk, indistinct and noisy (fig. 4A3-B2). In contrast, average CDS part *GC*3 gradients of both species are clearly distinct among classes, more regular and enlarged compared to the corresponding species specific average CDS part *G* + *C* gradients (fig. A4-B3). Finally, unlike all the other gradients which are U-shaped, *A. thaliana* average CDS part *GC*1 gradients tend to be bell-shaped. All the gradients are consistent among classes suggesting the existence of coordinated changes among codon positions (more distinct gradients can be seen in supplementary figures S7-S9).

In each species as expected, introns are *G* + *C*-poor compared to CDS parts (fig. 4A2-B1). Like CDS parts, average intron *G* + *C* content varies with both intron number and intron rank in the two species. In *A. thaliana* (fig. 4A2), for low intron number classes, average intron *G* + *C* content is mainly determined by intron number, firstly increasing with intron number for gene classes with few introns before stabilizing at *G* + *C* content around 32 – 33% for gene classes with more than five introns. In addition, a slight tendency towards an increase with intron rank along the genes is observed for low intron number classes. In rice (fig. 4B1), distinct and regular U-shaped gradients modulated by intron number are observed. Like for CDS part gradients, they are truncated for low intron number and otherwise highly regular albeit the amplitudes of the gradients are small compared to rice CDS part gradients. In this species, first intron *G* + *C* contents show a distinct increase with intron number while last introns stabilize at 45% of *G* + *C*, *G* + *C* content in the lower part of the U stabilizing above 33%. Like for CDS part gradients albeit changes in *G* + *C* content are less large, differences among genes within genomes and between the two genomes tend to decrease with the increase in intron number and the rank of the introns. Indeed, internal introns of genes with a high intron number tend to reach similar *G* + *C* content around 32%, the bottoms of the rice U-shaped gradients decreasing towards this value while *Arabidopsis* introns tend to increase towards it.

To sum up, large variation in *G* + *C* content intricately associated with intron location are observed in intronic genes, introns being not only *G* + *C*-poor but appearing to structure coding regions in step-gradients that vary in amplitude and shape according to the level of organization studied. In addition, gradients are also observed in introns. In contrast, in intronless genes gradients are smaller.

### Intron insertion within 5’ or 3’ UTRs modifies intron and CDS part *G* + *C* content in directions predicted by gradients

To test for the implication of introns in shaping CDS part *G* + *C* content, we compared genes having a single intron inserted either into the 5’ or the 3’ UTR with genes having the same number of introns inserted within their CDS and no intron inserted within UTRs. In both species for all intron numbers studied (up to eight introns inserted within CDS), intron insertion in 5’ respectively 3’ UTRs modifies *G* +*C* content of downstream respectively upstream sequences in both introns and CDS parts located dozens to thousand nts away from the insertion sites in a predictable direction that depends on the gradients described in the two species.

Regarding CDS part *G* +*C* content, the effect of intron insertion in UTR is illustrated in fig. 5AB for genes with seven introns inserted within CDS (see supplementary figure S10 for other number of introns). In *A. thaliana* and rice, intron presence in 5’ UTR is associated with a significant decrease in *G* + *C* content of the first CDS part that leads it to a *G* + *C* content roughly similar to a second CDS part of genes having no intron inserted within UTRs. In rice, a systematic decrease in *G* + *C* content is also observed when an intron is present in 3’ UTR while in *A. thaliana*, although decreases are generally observed they are not always significant. This pattern is consistent for all intron number classes with low intron numbers. In rice, intron insertion in 5’ or 3’ UTR also affects patterns of *G* + *C* content in *GC*1, *GC*2 and *GC*3 in the direction predicted by the gradients observed in genes with no intron insertion within UTRs (supplementary figures S11-S13). In *A. thaliana*, systematic significant differences are observed only in *GC*3 and not in *GC*1 and *GC*2 (supplementary figures S11-S13). In contrast, no systematic decreases are observed in the extremities of the genes opposed to intron insertion indicating that the location of the intron is determinant. Tests of significance are provided in Supplementary tables S5-S12.

**Figure 5:**
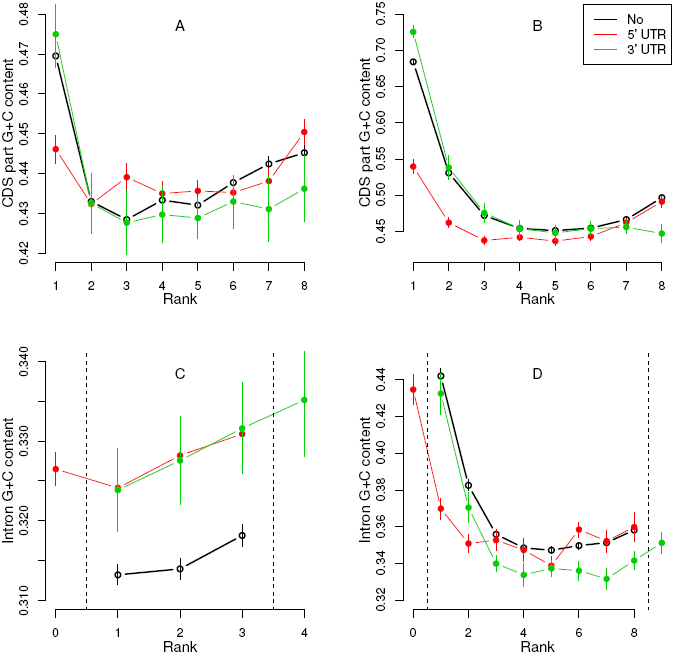
Comparison between genes having an intron inserted within one UTR with genes having no intron inserted in UTRs. In each cases, the comparison is made between genes having the same number of introns within coding regions and differing by the presence or the absence of an intron in their UTRs (no intron present in UTR: black, an intron in 5’ UTR: red, an intron in 3’ UTR: green). A. Average CDS part *G*+*C* gradients of genes having seven introns inserted within CDS in *A. thaliana*. Intron insertion in 5’ UTR leads to a decrease in first CDS part *G* + *C* content. B. Average CDS part *G* + *C* gradients of genes having seven introns inserted within CDS in *O. sativa*. Intron insertion in 5’ or 3’ UTR leads to a corresponding decrease in first or last CDS part *G* + *C* content. The dotted lines in panel C and D delimit the coding regions. C. Average intron *G* + *C* gradients of genes having two introns inserted within CDS in *A. thaliana*. Intron insertion within the 5’ or the 3’ UTRs leads to an increase in the *G* + *C* content of all downstream introns or upstream introns. D. Average intron *G* + *C* gradients of genes having seven introns inserted within CDS in *O. sativa*. The additional intron is integrated within the gradient as first (when inserted within the 5’ UTR) or last (when inserted in the 3’ UTR) and leads to a shift along the gradient of the downstream or the upstream introns inserted within the coding regions. Bars on dots represent standard error of means.

Regarding intron *G*+*C* content, the effect of intron insertion in UTR is illustrated in figure 5C for genes with three introns inserted in CDS for *A. thaliana* and figure 5D for genes with seven introns inserted in CDS in rice (see supplementary figure S10 for other number of introns). In *A. thaliana*, the increase in gene intron number is associated with an overall increase in *G* + *C* content of all the introns. As shown in figure 5C for genes with three introns inserted within CDS, the presence of an intron in both UTRs also lead to an overall increase in *G* + *C* content of all the introns. This pattern is consistent for all intron number classes with low intron numbers. In rice, the UTR’s intron is added to the gradient at the relevant extremity while the next intron (the second when the intron is inserted in the 5’ UTR or the penultimate when the intron is located in the 3’ UTR) presents in all cases a significant decrease in *G* + *C* content that drives then at a *G* + *C* level similar to a second or a last CDS part introns of genes with no intron inserted in their UTRs. Again, this pattern is consistent for all intron number classes with low intron numbers.

Similar results are obtained in each species with a subset of pairs of paralog genes differing by the presence of an intron in the 5’ or 3’ UTR of one member of each pair, both members of the pairs having the same number of introns inserted within coding regions. Although no further information on the time of gene duplications, the time or the types of events (deletion of the UTR intron in one of the duplicate or insertion of one intron in one of the duplicate) were used, whenever paired Wilcoxon signed-rank tests indicated the existence of significant differences, they were in the same direction of those described above in the two species confirming the hypothesis that changes in intron structure are implicated in the changes in *G* + *C* content (see supplementary tables S13 and S14).

To sum up, intron insertion in 5’ respectively 3’ UTRs modify *G* + *C* content of downstream respectively upstream sequences in a different way according to the types of regions (intron or coding regions) or the codon positions. The additional intron is integrated into the intron gradient while the CDS part gradient is shifted as expected if the intron inserted in UTR had the same effect as an intron inserted within the CDS on downstream coding regions.

### Intron has a specific impact on nt composition of coding regions

Introns located dozens to hundreds of nts away can affect *G* + *C* content in both introns and coding regions. To test if this effect is purely due to the addition of a given number of nts that takes away coding regions from transcription start sites (TSS) or if there is a specific intron effect, we performed two different kinds of comparisons. Firstly, we compared first CDS part *G* + *C* content between genes having no intron inserted within their 5’ UTRs and genes having an intron inserted within their 5’ UTR’s and similar distances between the TSS and the start codon. When no intron is inserted within 5’ UTR this distance is equal to 5’ UTR length while when an intron is inserted, the distance is equal to the sum of the two 5’ UTR part lengths plus the length of the intron. Indeed, a significant decrease in first CDS part *G* + *C* content is observed in both species when an intron is present within the 5’ UTR compared to pure 5’ UTR of comparable length suggesting that introns have a larger impact than 5’ UTR sequences (fig. 6AB). Secondly, in the subsample of genes with no intron inserted within UTRs we sorted genes into two classes according to the length of their first introns (shorter or longer than 150 nts in *A. thaliana* and 250 nts in rice, see supplementary figures S14 and S15 for additional information) and compared *G* + *C* content of the second CDS part in genes having a similar distance between TSS and the beginning of the second CDS part (this distance is equal to the sum of the 5’ UTR, first CDS part and first intron lengths). Again, an intron specific effect is detected in both species (fig. 6CD). In rice, the effect of first intron length on second CDS part *G* + *C* content takes the form of a huge decrease in second CDS part *G* + *C* content which appears to depend in a threshold way on the length of the first intron and can even affect all downstream CDS parts or introns in genes with few introns (for additional information see supplementary fig. S15). In *A. thaliana*, a weak albeit significant increase of second CDS part *G* + *C* content is observed between short and long introns (fig. 6C). All these results suggest that intron effect on coding region *G* + *C* content is specific and differs from other types of regions (5’ UTR or coding region).

**Figure 6:**
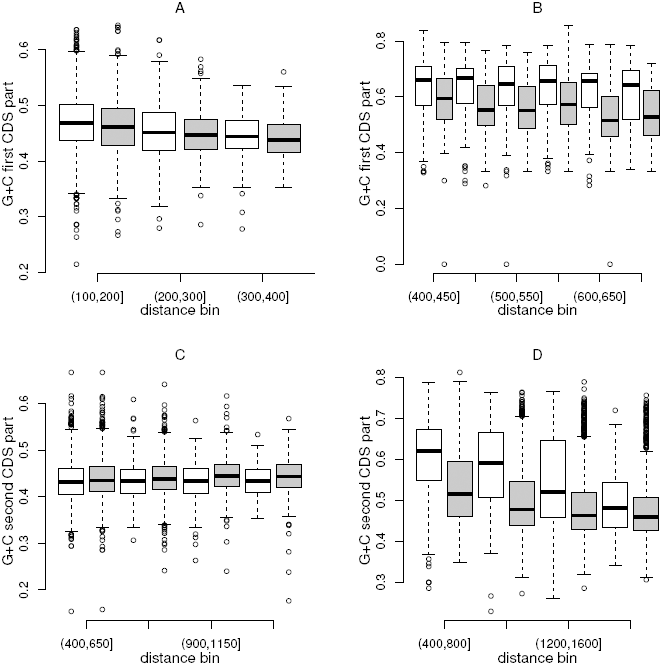
Intron specific effect. AB. Comparison between genes differing by the presence/absence of an intron within the 5’ UTR. Six bins of distance between the transcription start site and the translation start sites were formed for genes with no intron in UTRs (white) and genes with a single intron in the 5’ UTR (grey). For each bins, the boxplots show the *G* + *C* content of the first CDS part and in both species a decrease is observed between UTR alone and UTR plus intron. A. *A. thaliana*: sign-ranks tests within bins were all significant (*p <* 0.05, in most cases *p <* 0.001) except for the two last bins. B. *O. sativa*: sign-ranks tests within bins were all significant (*p <* 0.0001 except for the last bin *p <* 0.01). CD. Intron length threshold effect. Genes with no intron inserted within UTR were sorted into two groups according to the length of their first intron (below 150 nts: white, above 150 nts grey) for *A. thaliana* and (below 250 nts: white and above 250 nts: grey) for *O. sativa*. For each bins, the boxplots show the *G* + *C* content of the second CDS part. C. *A. thaliana*: an increase is observed in genes with long introns, sign-ranks tests within bins being all significant (*p <* 0.005). *B. O. sativa*: a decrease is observed in genes with long introns, sign-ranks tests within bins being all significant (*p <* 0.0001 except for the last bin *p <* 0.05).

### Gradients along genes explain part of the genome-wide variation in CDS *G* + *C* content in rice and *Arabidopsis*

Previous sections indicate that introns are associated with the formation of step gradients modulated by intron number and as a result, one can expect that CDS *G* + *C* content varies with intron number within each genome. Indeed in both species as shown in figure 7AB, CDS *G* + *C* content distributions both shift to lower values and decrease in dispersions with increasing intron number. In rice, low intron number classes (zero to two introns) are multimodal, spanning the genomic range of CDS *G* + *C* content variation with major modes located at high *G* + *C* content. As intron number increases, the high *G* + *C* mode decreases in favor of lower *G* + *C* content modes and the distributions eventually become mainly unimodal from seven intron number (with a minor mode at intermediate *G* + *C* content) and after further shift and reduction in dispersion stabilize within the 40 − 55% range. In *A. thaliana* like in rice, (i) CDS *G* + *C* content of genes with low intron number span the whole genomic range of variation, (ii) CDS *G* + *C* content decreases in both location and dispersion with the increase in intron number and (iii) for high intron numbers (above 10 introns) distributions appear to stabilize at low *G* + *C* content in a narrow range (40 – 49%).

**Figure 7:**
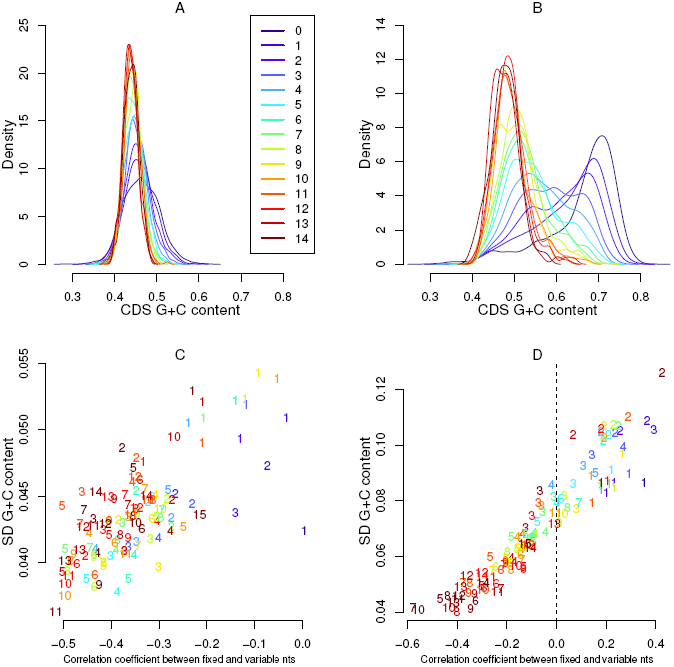
CDS *G* + *C* content distribution according to intron number in *A. thaliana* and *O. sativa*. Gene intron number is indicated in all panels by the colors (legend in panel A). AB. density outlines of CDS *G* + *C* content according to intron number. A. *A. thaliana* densities mainly decrease in width with increasing intron number while their modes shift slightly towards lower *G* + *C* content. B. In *O. sativa*, low intron number densities are multimodal with a *G* + *C*-rich major mode that decreases in height as intron number increases in favor of the *G* + *C*-poor lower modes to eventually become unimodal and *G* + *C*-poor from nine introns onward. Rice bimodality is largely due to the combined decreases in CDS *G*+*C* content and gene counts with intron number. CD. *G*+*C* content standard deviation (SD) within CDS parts *versus* correlation among non-synonymous (fixed) and synonymous (variable) sites. Points were replaced by number indicating the CDS part rank, the colors indicating the gene intron number. In both species, as intron number increases a decrease in correlations coefficients and SD is observed within CDS parts of a given rank. The decrease in both correlation coefficient and SD is more important in internal CDS parts. In *A. thaliana*, all correlation coefficients are negative while in rice, according to intron number and rank they vary from positive (in genes with low intron numbers or in external CDS parts of genes with high intron numbers) to negative (in central regions of genes with high intron numbers).

To gauge the link between gradients and CDS *G* + *C* content distributions, we regressed gradient amplitudes within intron number classes (equation 8) against *G* + *C* content medians of the classes. As shown in table 1, the ability of gradient amplitudes to predict changes in CDS median with intron number is fairly high in *A. thaliana* and can hardly be higher in rice for both codon *G* + *C* content and *GC*3. It is moderate in *A. thaliana* to high in rice for the two other codon positions. Furthermore as expected if gradients were involved in whole CDS patterns of variation, with the increase in gene intron number a decrease in CDS *G* + *C* content is observed for all decreasing gradients while the bell-shaped gradients of first codon position in *A. thaliana* are associated with an increase of CDS *GC*1 median with the increase in intron number (for additional information see supplementary Table S3 and S4 and supplementary figures S7-S9). Hence in both species, step gradients and their consequences on CDS *G* + *C* content permit to explain part of the genome-wide distribution in CDS *G* + *C* content, especially in rice where the bimodality of the overall distribution (fig. 2B) is largely due to the combined decrease of CDS *G* + *C* content and gene counts with intron number.

**Table 1.** Relationships between intron number class gradients and whole CDS *G* + *C* content in *A. thaliana* and *O. sativa*. *M*_*i*_ is the median *G* + *C* content or codon position *G* + *C* content (first position *GC*1, second *GC*2 and third *GC*3) of whole CDS of genes and *G*_*i*_ the respective gradient amplitudes within an intron number class, *i ∈ {*1*, …,* 14*}*.

| Type | *A. thaliana* | *O. sativa* |
| --- | --- | --- |
| *GC1* | $M_i = 0.5 + 0.41G_i, R^2 = 0.37, p = 0.02$ | $M_i = 0.66 - 0.77G_i, R^2 = 0.95, p < 10^{-4}$ |
| *GC2* | $M_i = 0.41 - 0.18G_i, R^2 = 0.39, p < 10^{-4}$ | $M_i = 0.51 - 0.55G_i, R^2 = 0.78, p < 10^{-4}$ |
| *GC3* | $M_i = 0.48 - 0.78G_i, R^2 = 0.89, p < 10^{-4}$ | $M_i = 0.72 - 0.97G_i, R^2 = 0.99, p < 10^{-4}$ |
| *G + C* | $M_i = 0.46 - 0.58G_i, R^2 = 0.88, p < 10^{-4}$ | $M_i = 0.72 - 0.91G_i, R^2 = 0.98, p < 10^{-4}$ |

### The decrease in CDS *G* + *C* content dispersion is due to several factors including changes in correlations among codon positions

Codon position gradients display coordinated patterns of variation along genes modulated by gene intron number. To study correlations according to site positions within CDS parts, we sorted coding nts into two groups depending on whether the nt can shift from *A/T ↔ G/C* without altering the amino acid sequence of the protein (synonymous nts, V) or not (non-synonymous nts, F) and computed Pearson correlation coefficients between *G* + *C* content of these two groups of nts within CDS part. In both species, a variation in the correlations that depends on the rank of the CDS part and gene intron number is observed (fig. 7CD). In *A. thaliana*, the correlations are negligible in genes with low intron number but become more and more negatives in internal CDS parts as gene intron number increases. In rice, they vary from largely positives in first and second CDS parts to largely negative in internal regions of genes with high intron numbers. The decomposition of CDS variance indicates that these changes in sign and value of correlations along genes contribute to roughly a one third decrease in CDS *G* + *C* content variances between genes with low and genes with high intron number in both species (see supplementary tables S15 and S16). While positive correlations between positions were already reported (e.g. Serres-Giardi et al., 2012), properly taking intron structure into account allowed revealing a complex pattern of variation in correlations along the genes including negative correlations, which were not previously documented, as far as we know. There are probably several other causes to the decrease in CDS *G* + *C* content variance with intron number (see supplementary fig. S16 for the effect of length variation on variances) and further investigations will be required to fully characterize the complex variance-covariance patterns of variation in *G* +*C* content along genes.

### *G* + *C* content variations both within and between genomes are much higher in external than in internal gene regions

Despite a strong difference in their overall *G* + *C* content, *A. thaliana* and rice exhibit common trends. Observations of CDS part and intron gradients along genes indicate that for both, differences within genome among intron number classes are huge in 5’ external CDS parts and introns and vanish in gene internal regions as intron number increases. Moreover, differences in internal gene regions of genes with high intron number do not vanish among genes within a genomes but also between genomes. For example when one compares CDS part and intron median gradients of genes with 11 introns in both species, gradients are all almost flat in the internal regions of genes. No differences in median *G* + *C* content can be detected for CDS parts of rank from five to 11 neither among ranks within a species nor between species while intron medians tend to be fairly close for intron ranks comprise between four and ten (fig. 8A). Likewise in coding regions, while each codon position reaches a given level of *G* + *C* content according to rank position along genes and intron number, as intron number increases the differences among internal CDS parts within a species or between species tend to disappear for each codon position (fig. 8B for genes with 11 introns). In contrast, differences among external CDS parts or introns and between species are large. In these regions, the gradients are steeper and different patterns of variation that are species specific and linked with the nature of the regions (coding regions or introns) or the codon positions are observed. To gauge the extent of overlap or divergence among regions within species and between species, we performed quantile-quantile plots of *G* + *C* content of CDS parts respectively introns of the same rank and having the same number of introns between the two species. For all intron number classes, for both coding regions and introns and within coding regions for each codon position, external CDS parts or introns are largely different while all *G* + *C* content distributions in internal regions overlap more and more as intron number increases (see fig. S17). Genes with few introns being mostly composed of external regions, they are always highly different between species. Hence, introns appears to delimit gene space in three regions, an internal region characterized by a conservation of *G* + *C* content levels among genes within a genome and between species and two external regions submitted to species-specific factors of varying intensities modifying *G* + *C* content which differ from internal regions within a genome and between the species.

**Figure 8:**
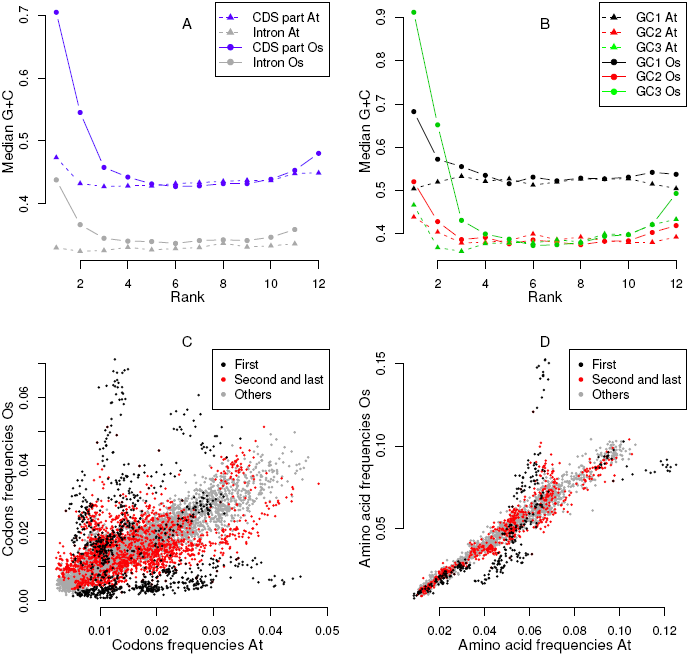
Comparisons of *G* + *C* content patterns of variation along genes within and between genomes. A. Gradients in median *G* + *C* content for CDS part (blue) and intron (grey) *G* + *C* content according to rank along genes in *A. thaliana* (triangle and dashed lines) and *O. sativa* (solid dots and plain lines). B. *GC*1 (first position within codon, black), *GC*2 (second position within codon, red), *GC*3 (third position within codon, green) gradients in median per codon position. All gradients differ in amplitude and in shape for each position within codons as well as for all nt positions within codons and introns. The central regions of gradients are strikingly close among rank along genes and between species, while external regions differ markedly especially in the 5’ regions. C. Codon frequency comparisons between *A. thaliana* and *O. sativa*. Each dot represents the frequency of a codon for a given CDS part and gene intron number in *A. thaliana* on the x-axis and *O. sativa* on the y-axis. D. Amino acid frequency comparisons between *A. thaliana* and *O. sativa*. Each dot represents the frequency of an amino acid for a given CDS part and gene intron number in *A. thaliana* on the x-axis and *O. sativa* on the y-axis. First CDS parts are plotted in black, second and last CDS parts in red and all other CDS parts are plotted in grey.

### Besides *G* + *C* content, codon and amino acid usage also vary with intron number and rank along genes

As *GC*1 and *GC*2 form gradients along the genes, one can expect to observe also variations in codon and amino acid frequencies linked with intron number and rank along genes. Indeed, there are few differences between *A. thaliana* and rice in internal regions of genes for codon (fig. 8C) or amino acid (fig. 8D) frequencies while a strong difference for both is evidenced at both ends of the genes. Codon and amino acid patterns of variation are consistent with *G* + *C* content patterns of variation (for additional information, see principal components analyses of both codon and amino acid frequencies, supplementary figure S18). Hence both codon and amino acid compositions are thus affected by intron number and vary according to CDS part rank along the gene in a genome dependent manner in external gene regions while internal gene regions tend to be similar even between species.

## DISCUSSION

In plant genes, patterns of variation in nt composition are highly complex and by the way difficult to describe. This might explain why previous studies failed to find the new information we provided here. Previous studies either neglected introns or did not account for them properly, focusing either on coding region *G* + *C* content (Wong et al., 2002), codon or amino acid frequencies (Wang et al., 2004; Wang and Roossinck, 2006; Wang and Hickey, 2007; Shi et al., 2007; Guo et al., 2007a; Mukhopadhyay et al., 2007), or investigating intron number and locations along genes separately (Zhu et al., 2009). Unlike previous studies, we undertook a systematic investigation of *G* + *C* content variation at different scales taking advantage of both the large number of genes and the wide variation in gene intron number characterizing plant genomes to address simultaneously the effect of gene intron number and intron location along the genes on *G* + *C* content for both coding regions and introns. In two widely different plant genomes, our analyses reveal that patterns of variation in *G* + *C* content are highly structured, more variable than previously described and that introns are both part of the patterns and implicated in their genesis.

In the two species, introns are intricately associated in the variation in *G* + *C* content at all of the scales investigated. Like in other eukaryotes, plant introns are *G* + *C*-poor sequences compared with coding regions that are *G* + *C*-rich, leading to the formation of conspicuous switch-back patterns in *G* + *C* content due to the alternation of CDS parts and introns along the genes. At nt scale, transitions between CDS part and introns are sharp, suggesting that widely different forces are at work to produce or to maintain the observed differences between coding regions and introns. Consecutive CDS parts display discrete changes in *G* + *C* content suggesting that there is no transcriptional unit with regard to *G* + *C* content but three different regions (two external regions and the internal region) delineated by the location of the introns. Intron insertion within 5’ or 3’ UTR decreases the CDS part gradient in the region of insertion and leaves it roughly unaltered in the other side of the gene. This indicates that CDS part *G* + *C* content is not determined by rank position with respect to coding regions but rather by the number of introns located upstream or downstream. Moreover, additional introns are included into a normal-looking intron gradients at the relevant extremity in rice while in *A. thaliana* it leads to an overall increase in all introns. As a result, intron insertion outside coding regions affect *G* + *C* content several dozens to hundreds of nts away in the same way as an intron located within the coding region. Further albeit limited supports of intron implication are provided by similar comparisons made this time within a set of related paralogs differing by the insertion of an intron in UTRs. In this subsample as well, introns shape *G* + *C* content, even if these last results are difficult to interpret because (i) the sample sizes of the different gene subsets are small and (ii) intron insertion in UTRs within paralogous genes are often associated with other changes of structure like changes in intron position or lengths and changes in CDS part lengths (Xu et al., 2012). All these results converge to support a major role for introns in shaping plant gene *G* + *C* content.

The decrease in *G* + *C* content associated with the presence of introns in UTRs suggests that introns might act as a barrier, preventing the overall increase in *G* + *C* content occurring in external gene regions to spread within internal gene regions (CDS parts or introns that are downstream and upstream at least one or more introns). An intron barrier effect is further supported by the observation that the intron effect is not reduced to a size effect. For a similar length and position from the TSS, introns have a larger impact than 5’ UTRs in both species or coding sequences in rice. The intron barrier effect hypothesis provides a simple explanation to many of the particularity of *G* + *C* content patterns of variation in the two studied species. It also probably applies to other plant species although further investigations will be required to confirm the generality of our observations. Indeed, in the context of such an intron barrier effect, U-shaped gradients appear as consequences of intron presence, while intron absence or low number of introns explain why genes with few introns are different from the others in gradient shapes and amplitudes. Furthermore, it could explain a part of the reduction of variance in CDS part *G* + *C* content which is observed with the increase in intron number. Finally, it might also explain why codon positions are affected in a coordinated way by introns and why gradients are modulated by intron number in a highly reproducible way among intron number classes, as a single mechanism might be responsible for all these patterns. Provided that indeed similar forces are shaping *G* + *C* content patterns of variation in plants, the combination of the intron barrier hypothesis and the species-specific factor determining the level of *G* + *C* content are sufficient to explain the sort of syndrome observed in plants of the combined variation in genome-wide *G* + *C* content, within genome among gene heterogeneity and gradient amplitudes.

Patterns of variation along genes can be interpreted as resulting from opposite forces shaping *G* + *C* content along genes. Within each genome, comparison of *G* + *C* content between genes with different intron numbers indicate that internal gene regions (CDS parts and introns that are located both downstream and upstream one or several introns) have similar *G* + *C* content distribution while external gene regions (first and second, penultimate and last CDS parts) are different from internal ones. A surprising result, which generality requires confirmation in other plant species, is that internal gene regions of the two studied species are in fact similar. On the opposite, patterns of variation in genes with low intron number or external gene regions exhibit species-specific differences with large difference in *G* + *C* content but also varying patterns of variation in introns and in coding regions according to codon positions. The existence of differences between external and internal regions of genes is further supported by the changes in the sign or the importance of correlations observed within CDS parts between the *G* + *C* content of nts that can change from *A/T ↔ G/C* without altering the amino acid sequence of the proteins (hereafter referred to as synonymous nt, all third codon positions plus first codon positions of Leucine and Arginine) and those that cannot (non-synonymous nts, all second codon positions and other first codon positions). To produce such a pattern of variation along the genes, conflicting forces are required. As already pointed, in external gene regions species-specific forces seem to be at work. As mentioned in the introduction and extensively described in Glémin et al. (2014), GC-biased gene conversion driven by recombination gradient could be involved to produce the species-specific increase in *G* + *C* content in external gene regions. If this hypothesis is correct, our results suggest that introns act as a barrier to recombination or to the extension of conversion tract through the middle of genes. In agreement with this hypothesis, recombination rates were found to be lower in introns than in CDS parts in *Mimulus guttatus* (Hellsten et al., 2013). *In internal gene regions, the raise of negative correlations among synonymous and non-synonymous nts suggests that constraints are applying on internal CDS parts to maintain G*+*C* content within a given range. Neither drift nor mutation are expected to produce negative correlations among codon positions and GC-biased gene conversion is expected to produce positive correlations because it affects all positions in the same direction. To our knowledge, stabilizing selection on *G* + *C* content appears as the only phenomenon able to produce these correlations. The possible cause of this stabilizing selection are unknown but it could be related to the maintenance of distinct *G* + *C* content between CDS parts and introns. For example, the differences in *G*+*C* content between introns and CDS parts are described as promoting intron recognition and splicing in *A. thaliana* (Goodall and Filipowicz, *1991*; Amit et al., 2012; Gelfman et al., 2013), in maize (Carle-Urioste et al., 1997; Ko et al., 1998; Clancy and Curtis, 2002) but also in other eukaryotes (Amit et al., 2012). At DNA sequence levels, nucleosome occupancy is tightly associated with *G + C* content (Tillo and Hughes, 2009) and more generally gene coding regions in eukaryotes including *A. thaliana* (Choi et al., 2013) *while introns tend to be linkers. Studies in yeast, fishes and nematodes suggest that nucleosome occupancy is associated with strong alterations of mutation biases (Chen et al., 2009, 2012) while in yeast, it has been suspected to lead to stabilizing selection in G* + *C* content (Kenigsberg et al., 2010) providing both a possible feed-back loop compatible with our observations and a possible explanation for the negative correlations among codon positions within CDS parts. Whether the constraints on *G* + *C* content within internal CDS parts could participate to the potential intron barrier effect remain to be investigated as well as other potential causes.

Introns are required to properly describe not only the patterns of variation in *G* + *C* content in protein-coding genes but also the variations in codon and amino acid frequencies. Beyonds their proper description how these patterns have been created and are maintained remains an open question that is unlikely to be solved without taking into account the fact that variation in *G* + *C* content is not only tightly associated with introns but also with variation in recombination rates, patterns of methylation and nucleosome occupancies, this list being probably non-exhaustive (Glémin et al., 2014). Indeed, the correlations among these different features and *G*+*C* content patterns of variation might result either from a direct influence of one of these on all the others or from co-evolutionary processes including potentially positive feed-back loops. More generally, all the processes involved in protein production are suspected to affect nt compositions at DNA sequence or mRNA levels depending on whether they are related to transcription, splicing processes or mRNA folding (Chamary et al., 2006; Parmley et al., 2007; Warnecke et al., 2009; Shabalina et al., 2013). However only speculations are possible for the moment, gene intron structure being almost always neglected. To our knowledge, none of the studies investigating nucleosome positioning (Andersson et al., 2009; Schwartz et al., 2009; Tilgner et al., 2009; Tillo and Hughes, 2009; Chodavarapu et al., 2010), the distribution of recombination rate within genomes (Hellsten et al., 2013; Choi et al., 2013), the interplay between methylation and *G* + *C* content (Takuno and Gaut, 2013; Gelfman et al., 2013), and more generally mutational biases or protein rates of evolution investigated in an exhaustive way how these features are affected simultaneously by intron number and intron location along the genes, or were conducted in *S. cerevisiae*, a species remarkable for its scarcity in introns. Even when splicing is the focus of the studies (Amit et al., 2012; Gelfman et al., 2013), intron number and location of CDS parts or introns along genes are not described in detail. Our results suggest that it could be a serious flaw for three reasons. Firstly, although not properly taken into account, introns are regularly mentioned in previous studies and whenever an intron issue is addressed, it has an impact on the patterns described (Schwartz et al., 2009; Hellsten et al., 2013; Choi et al., 2013). Secondly, our own observations restricted to sequence *G* + *C* content demonstrate the existence of a strong structure associated with intron cutting up. Ignoring this structure leads at least to underestimate the breadth of the variation and can at worth be misleading. Indeed, gene intron number distribution is severely unbalanced and as a result, the observation made neglecting intron number are always biased in one way or another. And thirdly, variation due to intron number and location along genes are so strong (at least in rice) that even if patterns in *G* + *C* content are only a consequence of other processes and no co-evolution is occurring, neglecting introns when trying to identify the mechanisms in cause in *G* + *C* content variation or features showing a tight correlation with *G* + *C* content is likely to fail to account for the structuring effect of introns. As a striking illustration of this point, the likely role of stabilizing selection on CDS part *G* + *C* content was detectable only by properly taking intron structure into account. This was previously missed because at the whole gene level most variation was explained by processes affecting external gene regions. In conclusion, to study gene architecture evolution introns have to be taken into account at least within plant genomes.

## METHODS

### Genomic data

Genomic data come from whole genome annotations of *A. thaliana* and *O. sativa* var. nipponbare. Two main gene divisions can be envisioned. Genes can be divided into coding and non-coding regions and in this case four types of regions can be described, three non-coding (5’ UTR, introns and 3’ UTR) and the coding regions (fig. 1). Alternatively, genes can be divided between introns and exons, the latter composing the mRNA. In this case, exons can be of formed either purely of UTR if an intron is inserted in the UTRs, by a mixture of UTR and coding regions (for example in external exons of genes with no intron inserted in UTRs or intronless genes) or purely by coding sequences for the most internal exons. Preliminary analyses indicated that within exons composed of a mixture of UTR and coding sequence (i) significant differences in average *G* + *C* content occurred between coding and non-coding regions (paired t-test within intron number classes, all p-value *<* 10^−4^), and (ii) significant but low correlations were found between coding and non-coding regions (in *Arabidopsis*, Pearson correlation coefficient 5’ UTR/cds1 of 0.15, 3’UTR/last CDS part of 0.05; in rice Pearson correlation coefficient 5’ UTR/cds1 of 0.35, 3’UTR/last CDS part of 0.13; all p-value *<* 10^−4^). for these reasons, we decided to study all four types of regions separately (5’ UTR, coding sequence, intron and 3’ UTR) and to use the term CDS part for coding region pieces and discard the term exon. A gene can then be composed of a number of introns plus a number of 5’ UTR parts (0 if no 5’ UTR is described, 1 if no intron is present in the 5’ UTR and *n* + 1 if *n* introns are located in the 5’ UTR region of the gene) plus a various number of CDS parts (from 1 if no intron is inserted in the coding regions to *n* + 1 if *n* introns are inserted in the CDS) plus a various number of 3’ UTR parts (0 if no 3’ UTR is described, 1 if no intron is present in the 3’ UTR and *n* + 1 if *n* introns are located in the 3’ UTR of the gene). Along a gene, each region was denoted according to its type (5’ UTR, coding region, intron or 3’ UTR) and numbered according to its rank from the 5’ end towards the 3’ end (fig. 1).

#### A. thaliana

We used the TAIR10 genome release avalaible at ftp://ftp.arabidopsis.org/. A single gene model was collected per locus/protein-coding gene, the one having the longest CDS and the highest confidence score. For each of the selected models, the confidence score, the chromosome, the gene start and stop positions on the chromosome, the number of introns, the start and stop of each of the different elements composing the gene were recorded. Only genes with a confidence score higher or equal to four were conserved.

#### O. sativa

We used the Release 6.1 of the MSU Rice Genome Annotation Project available at ftp://ftp.plantbiology.msu.edu//. No confidence score similar to the one existing in *Arabidopsis* is provided. However, when expression data like ESTs or FLcDNA are available, it is indicated. We took only gene models supported by full length cDNA. For each of the selected gene models, the chromosome, the gene start and stop positions in nt along the chromosome, the number of introns, the start and stop of each of the different elements composing the gene were recorded.

#### Final datasets

We restricted our analyses to gene models which have both UTRs described and are supported by expression data ending up with a similar number of genes in both species (18134 in *A. thaliana* and 17862 in rice). We also removed all gene models for which at least one element was less than ten nts long to avoid to compute *G* + *C* proportions on too few nts. To avoid confusion about intron number, we decided to further restrict our analyses to three different sets of genes. In the two studied species, most of the genes do not have intron inserted in UTRs (around 79% in *A. thaliana* and 75% in rice). In these two subsets of genes, the distribution of intron number within coding regions is very similar (supplementary Table S1) and we decided to use these two subsets of genes as reference sets to describe the within genome among genes effects of intron number. In addition, in each species, we formed two additional datasets to test for an intron effect on nt composition: a first one made of the genes having any number of introns inserted within their coding regions plus a single intron inserted within their 5’ UTR, and for the second genes having any number of introns inserted within their coding regions plus a single intron inserted within the 3’ UTR (supplementary Table S1). While the 5’ UTR datasets comprise roughly the same number of genes in the two species, intron insertions in 3’ UTRs of rice are more than twice as frequent as in *A. thaliana*. In both species, the remaining set of genes (4024 in *A. thaliana* and 4438 in rice) display a high number of different combinations of intron insertion within UTRs, each represented only by a few number of genes and were discarded. Because of the low sample size, we also discarded genes with more than 14 introns in the two reference sets. The final composition of the six datasets is shown in supplementary Table S1. Finally, to further study the consequences of the insertion of introns within UTRs, we also retrieved the orthologs set of genes available on the MSU site at http://rice.plantbiology.msu.edu/annotation_pseudo_ortho.shtml. This dataset contains for both species the genes that are orthologous within a genome (duplicated genes) and we used it to produce in each species an additional dataset composed of duplicated genes that differ by the presence/absence of a single additional intron inserted either in the 5’ UTR or 3’ UTR (Supplementary table S2).

### Compositional analyses

#### *G* + *C* content measurements

In each species, genes were distributed into classes according to their intron number. Let *i* be the intron number of the genes and *N*_*i*_ the number of genes belonging to the *i*th class (*i* ∈ {0*, …,* 14}). Within classes, genes were denoted by the indices *g* (*g* ∈ {1*, …, N_i_*}). In each species for each of the selected gene model, gene sequences were retrieved and analysed using R and the bioconductor package (R Core Team, 2013; Gentleman et al., 2004). After having sorted genes according to intron number, we explored *G* + *C* content pattern of variation at different scales.

At nt level, to study transitions between coding regions and introns we computed *G* + *C* content on 50 nts spanning both sides of each exon-intron junction, sequences being aligned on the consensus 5’ and 3’ splice sites. With such an alignment, information of nucleotide position according to the Open Reading Frame (ORF) of the sequences are lost in coding regions. Each junctions were studied separately within intron number classes. Large differences being observed between introns and coding regions, to study patterns of variation within coding regions, we removed intron sequences and concatenated CDS parts, keeping information on CDS part rank. All coding sequences within an intron number class were then aligned according to their starting methionine (conserving ORF phase). Let *x*_*igl*_ be the *l*th nt of the *g*th gene with *i* introns and coding region length equal to *L*_*ig*_, *l* ∈ {1*, …, L_ig_*}. Patterns of variation in *G* + *C* at nt level were studied without taking into account intron cutting up of the CDS by simply computing

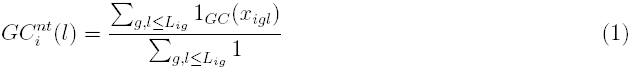

where 1_*GC*_(*x*_*igl*_) = 1 if *x*_*igl*_ ∈ {*G, C*} and 1_*GC*_(*x*_*igl*_) = 0 otherwise. Intron cutting up were took into account by computing *G* + *C* content per nt position according to the CDS part the nt belongs to

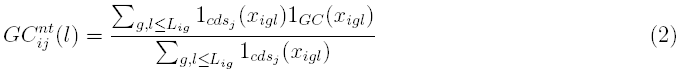

where 1_*cds*_*j*__ (*x*_*igl*_) = 1 if *x*_*igl*_ belongs to the *j*th CDS part and 1_*cds*_*j*__ (*x*_*igl*_) = 0 otherwise. *GC*1, *GC*2 and *GC*3 patterns of changes at nt levels are provided by sub-setting over the respective codon position in the former distribution 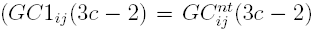, *GC*2_*ij*_(3*c* – 1) and *GC*3_*ij*_(3*c*), with *c* ∈ {1*, …, L_ig_/*3}). Codon *G* + *C* content is simply the mean of the three positions within a codon and were displayed at the second codon position leading to

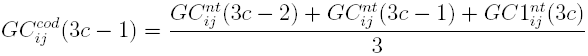

When a codon was interrupted by an intron, it was discarded from the computations. In all cases, *G* + *C* content were computed only when a nucleotide position were represented by at least 50 different sequences.

At element level (CDS parts or introns), *G* + *C* content of the different CDS parts or introns composing the genes were computed as the count of *G* and *C* nts over the total number of nts of the element and denoted 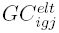 where *elt* indicates the type of the element (*cds* for CDS part, *I* for intron), *j* indicating the rank of the element in the gene decomposition (see figure 1). In addition in coding sequences, we computed the CDS part *G* + *C* content per codon position (denoted *GC*1_*igj*_ for the first, *GC*2_*igj*_ for the second and *GC*3_*igj*_ for the third position within codons) as the count of *G* and *C* nts at the given codon position over the count of nts at this codon position within the element. Average CDS part (or intron) *G* + *C* content were computed by averaging over all genes within an intron number class according to CDS part (or intron) rank, as shown in equation 4 for CDS part

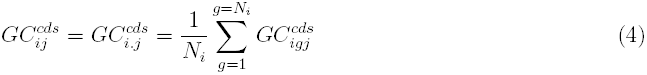

Intron cutting up were compared with a bin division where CDS were divided into the same number of equal-sized bins as the number of CDS parts (equal to the number of introns plus one). For a gene *g* with *i* introns, when CDS length were not a multiple of *i* + 1, all bin lengths were equal to └*L*_*ig*_/(*i* + 1)┘ except for the last one which were composed of the rest of the sequence. *G* + *C* content were computed in bins as done for intron cutting up.

#### Codon and amino acid analyses

For each of the two species, in addition to coding sequence *G*+*C* content analyses, we also computed CDS part codon and amino acid frequencies within classes. Stop codons as well as codons or amino acids overlapping between CDS parts were discarded. Let *nc*_*igj*_ be the vector of the counts of the 61 codons within the *g*th CDS of the *j*th CDS part of the genes with *i* introns, codon frequencies were computed as

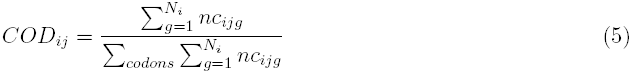

where *COD*_*ij*_ referred to the vector of frequencies of the 61 non-stop codons observed at the *j*th rank in genes with *i* introns. Let *naa*_*igj*_ be the vector of the counts of the 20 amino acids (aa) within the *g*th CDS of the *j*th CDS part of the genes with *i* introns, amino acid frequencies were computed as

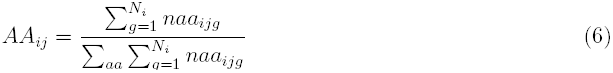

where *AA*_*ij*_ referred to any of the 20 amino acids observed at the *j*th rank in genes with *i* introns.

### Intra-species comparisons between genes with or without intron inserted within UTR

In each species, we compared the reference dataset composed of genes with no intron inserted in UTRs with each of the two datasets composed of genes having either an intron inserted in their 5’ UTR or in their 3’ UTR. Comparisons between the data sets were made between genes having the same number of introns inserted within their coding regions, the two datasets differing by the presence/absence of a single intron located either within the 5’ or 3’ UTRs. Within the different intron number classes, we made Welsh-two sample student tests on first resp. last CDS part to test for the existence of significant differences in *G* + *C* content. In addition, to investigate for an intron specific effect, we took both the 5’ UTR datasets and the intron-free UTR datasets and sorted genes into bins according to the distance between the transcription start site and the translation start site. Within each bin, we then tested for differences in *G* + *C* content between the two sets of genes with a sign-rank test (Wilcoxon test). We made the same kind of analyses on the intron free UTR gene subsets, sorting genes into two classes (short *versus* long first intron, below and above 149 and 245 nts respectively in *A. thaliana* and rice) and then in bins according to the distance between the transcription start site and the beginning of the second exon. In all cases, gene counts within bins were higher than 25 genes and in most cases higher than a hundred. Finally, in the subset of paralog genes, we formed pairs of duplicated genes composed of the gene without intron inserted within UTRs and gene with an intron inserted within either the 5’ or the 3’ UTRs. Genes from different intron number classes were pooled and Wilcoxson-paired tests were performed on respectively second CDS part, respectively penultimate CDS part *G* + *C* content and codon position *G* + *C* content for genes having a additional intron inserted within their 5’ respectively 3’ UTRs. In the same way, Wilcoxson-paired tests were performed on respectively first intron inserted in coding regions, respectively last intron inserted in coding regions *G* + *C* content for genes having an additional intron inserted within their 5’ respectively 3’ UTRs. In all cases, a Bonferroni correction for multiple-comparisons was also performed.

### Intra-species relationships between CDS part and whole CDS *G* + *C* content

Whole CDS *G* + *C* content of a gene is the mean of its CDS part *G* + *C* content weighted by the proportion of the different CDS parts within the gene :

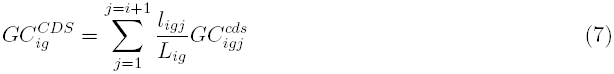

where *l*_*igj*_ is the length of the *j*th CDS part (*j ∈ {*1*, …, i* + 1*}*) and 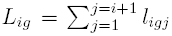 is the total number of nts of the CDS of the gene of interest.

As a result by construction, a link between *G* + *C* content gradients along genes and CDS *G* + *C* is expected. To gauge the links between intron number, patterns of variation along genes and whole CDS *G* + *C* content variation with intron number, we measured the average amplitude of the gradients along CDS within intron number class *i*, *G*_*i*_, for genes with one to 14 introns (intronless genes do not have CDS part gradients) :

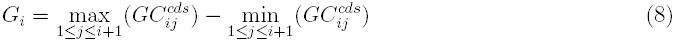

where 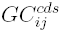 stand for the average CDS part *G* + *C* content of rank *j*. By construction, gradient amplitude is always positive independently of the shape of the gradient. Gradients amplitude according to codon positions were computed in the same way. We then regressed the median of CDS *G* + *C* content of the *i*th class against the gradient measured in the class against the whole CDS *G* + *C* content median of the class for the three codon positions (*GC*1, *GC*2 and *GC*3) and the codon (average over all three codon positions) *G* + *C* content within each species.

In addition, we investigated the patterns of variation in variance in relationship with intron number. Variances in CDS *G* + *C* content within intron number classes result from the balance between CDS parts variance in lengths and *G* + *C* content and on the covariances that occur between lengths, *G* + *C* content and the CDS parts along the genes. CDS part variance can be further divided according to the nature of the nts (non-synonymous or synonymous). The non-synonymous nts (denoted *F* ) cannot shift either from A or T to G or C or the converse without altering the amino acid sequence of the protein and the synonymous ones (denoted *V* ) are those nts for which *AT ↔ GC* mutation is possible without altering the amino acid sequence. If *V*_*ij*_ and *F*_*ij*_ represent the respective contributions of CDS part synonymous and non-synonymous nts to CDS part *j G* + *C* content of genes with *i* introns, then

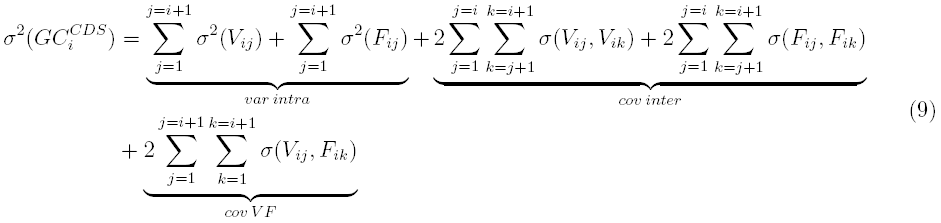

with 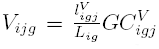 and 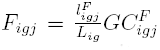, where 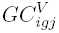 and 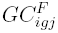, represent the *G + C* contents at synonymous and non-synonymous sites of CDS part *j* of the *g*th gene in the *i*th intron number classes and 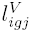 and 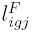 the number of each types of sites 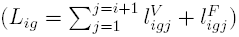.

### Inter-species comparisons of CDS parts *G* + *C* content, codon and amino acid frequencies

Inter-species comparisons of CDS part *G* + *C* content were performed on the two subsets of genes with no intron inserted within UTRs (reference sets) using quantile-quantile plots of CDS part *G* + *C* content distribution per rank within intron number classes. For codon and amino acid frequencies, principal component analyses were performed on the same datasets with codon or amino acid frequencies as the variables and CDS parts as individuals (each CDS part were identified by a “species”/”intron number”/”rank”) for the two reference sets of genes.

